# A systematic genetic interaction map of human solute carriers assigns a role to SLC25A51/MCART1 in mitochondrial NAD uptake

**DOI:** 10.1101/2020.08.31.275818

**Authors:** Enrico Girardi, Giuseppe Fiume, Ulrich Goldmann, Celine Sin, Felix Müller, Sabrina Lindinger, Vitaly Sedlyarov, Ismet Srndic, Benedikt Agerer, Felix Kartnig, Eva Meixner, Anna Skucha, Manuele Rebsamen, Andreas Bergthaler, Jörg Menche, Giulio Superti-Furga

**Author notes:** Corresponding author: Giulio Superti-Furga, CeMM Research Center for Molecular Medicine of the Austrian Academy of Sciences, Lazarettgasse 14, AKH BT25.3, 1090 Vienna, Austria.

## Abstract

Solute Carriers (SLCs) represent the largest family of human transporter proteins, consisting of more than 400 members^1,2^. Despite the importance of these proteins in determining metabolic states and adaptation to environmental changes, a large proportion of them is still orphan and lacks associated substrates^1,3,4^. Here we describe a systematic mapping of genetic interactions among SLCs in human cells. Network-based identification of correlated genetic interaction profile neighborhoods resulted in initial functional assignments to dozens of previously uncharacterized SLCs. Focused validation identified SLC25A51/MCART1 as the SLC enabling mitochondrial import of NAD(H). This functional interaction map of the human transportome offers a route for systematic integration of transporter function with metabolism and provides a blueprint for elucidation of the dark genome by biochemical and functional categories.

---

Transmembrane transporters are key contributors to the energetic and metabolic needs of a cell. The largest human family of transporters is composed of Solute Carriers (SLCs), a diverse set of transmembrane proteins consisting of more than 400 members^2^. While several prominent members of the family have been the focus of extensive research, a large proportion of them remains uncharacterized^1^. Genetic interactions offer a powerful way to infer gene function by a guilt-by-association principle^5,6^, with negatively interacting pairs often sharing functional overlap. Positive interactions, on the other hand, can reflect regulatory connections or participation in the same protein complex. Fundamental studies in model organisms have systematically expanded our understanding of genetic regulatory networks and functionally annotated orphan genes^6,7^ and recent technological development, in particular the development of CRISPR-based approaches, made now possible to tackle systematic mapping of genetic interaction landscapes in human cells^8–12^.

To gain insight into the interplay and functional redundancy among SLCs, we characterized their genetic interaction landscape in the human cell line HAP1. We combined a panel of isogenic cell lines each lacking one of 141 highly expressed, non-essential SLCs^13^ (Fig. 1a, Extended Data Fig. 1a) with a CRISPR/Cas9 library targeting 390 human *SLC* genes as well as a set of 20 genes previously shown to be important for optimal HAP1 growth^14,15^ (Fig. 1b). Cellular fitness was employed as a readout as it integrates cellular physiology, is functionally informative and is readily measurable^5^. Genetic interactions were tested in full media as it allowed us to directly correlate our results with other extensive datasets available for this cellular system^13–15^. To determine the ability of our approach to identify functionally relevant interactions, we initially screened two non-essential metabolic genes (*CPS1, DBT*) with well-characterized biological functions^16,17^. Scoring of genetic interactions was achieved by implementing a gene-level linear model using DESeq2^18^ and yielded non-overlapping patterns of gene interactions (Fig. 1c, Extended Data Fig. 1b-c). We infected HAP1 wild type (wt) and HAP1 cells lacking dihydrolipoamide branched chain transacylase E2 (encoded by *DBT*), a member of branched-chain alpha-keto acid dehydrogenase complex (BCKD) responsible for the mitochondrial breakdown of isoleucine, leucine and valine, with our SLC-focused CRISPR/Cas9 library and collected samples at 2, 9 and 16 days post-infection (p.i.). Analysis of the genetic interaction profile 16 days p.i. showed that BCKD-deficient cells are sensitive to loss of amino acid transporters such as the glutamine transporters SLC1A5 and SLC38A5 as well as members of the SLC7 family (SLC7A4, SLC7A11, Extended Data Fig. 1b). Loss of BCKD, resulting in maple syrup urine disease (MSUD), is known to induce BCAA accumulation as well as cellular depletion of several amino acids including glutamate, glutamine and alanine^19^, highlighting the potential of our approach to connect a given gene with specific metabolic processes, in this case amino acid metabolism. Screening for SLC genes interacting with *CPS1* (encoding carbamoyl-phosphate synthase 1, a mitochondrial enzyme catalyzing the first step of the urea cycle) identified strong negative interactions with the bicarbonate transporter *SLC26A3*, the carnitine transporter *SLC16A9*, the orphan transporter *SLC35A5* and the mitochondrial transporters *SLC25A36* (pyrimidines) and *SLC25A44* (BCAA, Extended Data Fig. 1c). It was previously reported that CPS1, which accepts ammonia derived from amino acid catabolism and bicarbonate as substrates, plays a role in maintaining the pyrimidine pool in cancer cells^20^. Loss of CPS1 results in hyperammonemia, which can be alleviated by carnitine supplementation^21^, generation of glutamine from BCAA or negative regulation of one carbon metabolism, producing ammonia as end product^22^, consistent with the observed buffering interactions involving the glycine transporters SLC6A18/19 and the folate transporter SLC19A1 (Extended Data Fig. 1c). Overall, these results suggest that genetic interactions involving SLCs can identify specific metabolites and metabolic processes related to a given gene, both in terms of substrates and biological functions.

**Fig. 1.**
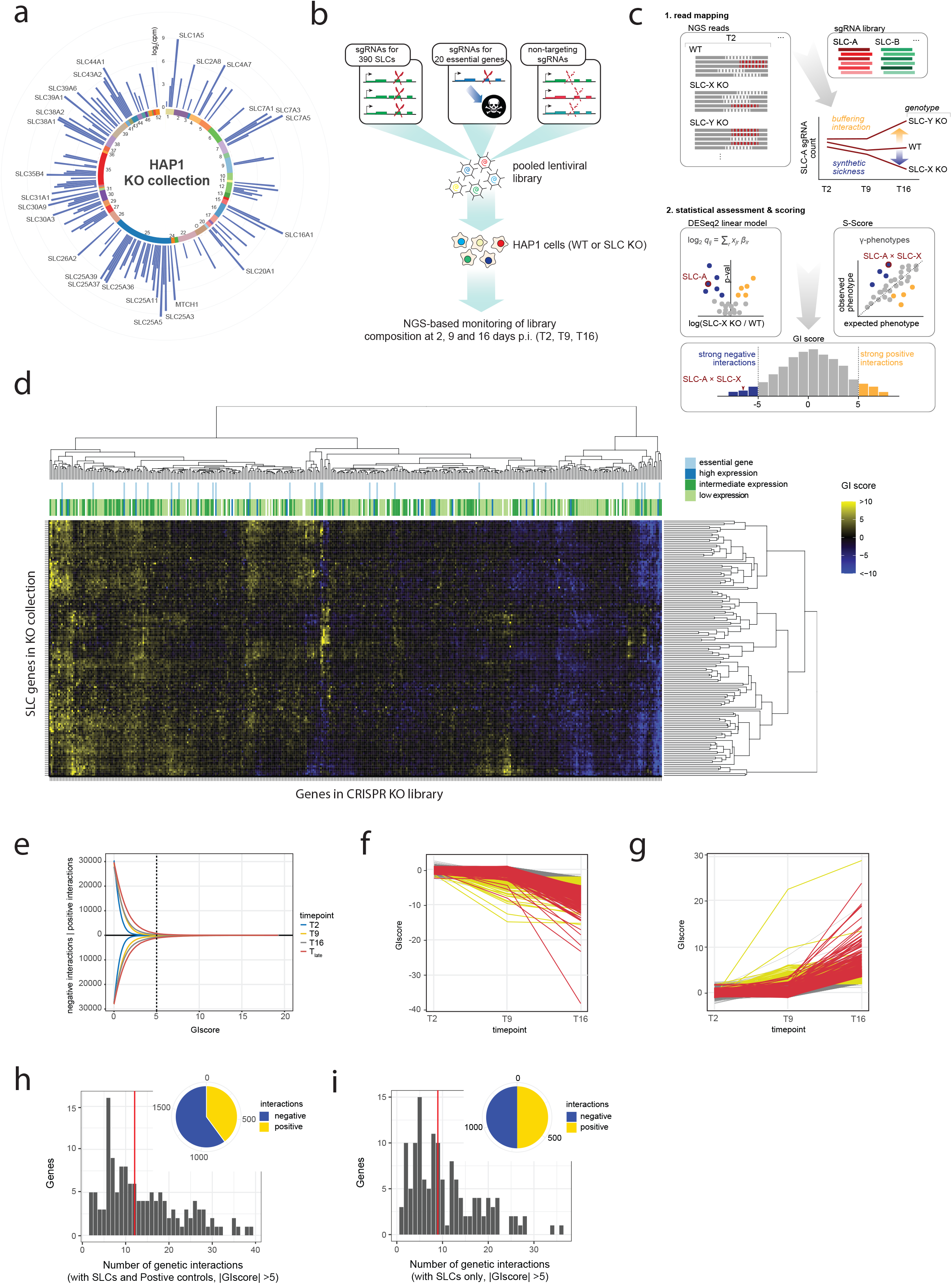
A genetic interaction landscape of Solute Carriers in human cells. (a) The panel of Solute Carriers genes deleted in HAP1 cells. Genes are arranged by subfamily, indicated by the number in the circle. Bars show the expression of the corresponding transcripts in wt HAP1 (count per million reads, cpm). Genes with expression above 2^6^ cpm are labeled. (b) Scheme of the composition of the CRISPR/Cas9 library used in the screen. (c) Scheme of the procedure to derive genetic interaction scores. (d) Heatmap of genetic interactions at the joint timepoint T_late_. Positive interactions are indicated in yellow, negative interactions in blue. Core essential genes in HAP1 cells according to^14^, as well as expression levels (low expression, cpm <10; intermediate expression, cpm between 10 and 100; high expression, cpm above 100) in HAP1 cells are indicated above the heatmap. (e) Number of genetic interactions depending on GI score threshold value and time point. The chosen cutoff for the subset of strong interactions is shown as a dashed line. (f-g) Time dependence of genetic interactions for the negative (f) or positive (g) subsets. “Fast” interactions are shown in yellow, “slow” interactions in red. (h-i) Histograms of the number of genetic interactions per gene within the set of all strong interactions (f) or SLC-SLC strong interactions (g). The median value is indicated in red. In the inset, the numbers of positive and negative interactions are shown.

To define the genetic interaction landscape of Solute Carriers, we extended this approach to our collection of SLC-deficient cell lines. Genetic interactions were derived using both DESeq2 and a modified version of the S-score previously used to quantify epistasis in yeast (Extended Data Fig. 1d-f)^23^. The two scores showed a robust correlation (Extended Data Fig. 1g) at the late time points and were linearly combined in a single score (named GI score) to identify the strongest interactions. We measured approximately 55.000 interactions per time point (Fig. 1c), observing a roughly equal number of positive and negative interactions as well as an increase in the strength of interactions over time (Fig. 1d-e, Extended Data Fig. 1g). Classification of interactions based on their sign and speed (Fig. 1f-g) showed that “fast” interactions involved genes expressed in HAP1 cells while essential SLCs were more likely to be involved in positive, rather than negative interactions (Extended Data Fig. 1h). To identify the most robust and time-independent interactions, an additional timepoint (T_late_) was also generated by merging the T9 and T16 timepoints. Strong genetic interactions are expected to be rare^5^ but the vast majority of the genes screened showed at least one strong (defined by an absolute GI score above 5) interaction with another gene present in the library (134/141, 95.0%, T_late_ timepoint), with a median of 12 interactions/gene (Fig. 1h). Selection of a set of strong SLC-SLC interactions (GI score above 5, 1414 gene-gene interactions, Fig. 1i) revealed a network of known and novel genetically interacting pairs (Fig. 2a, Extended Data Fig. 3a, interactive version available at http://sigil.cemm.at). In particular, we identified reciprocal negative interactions between functionally redundant paralogs, such as the two monocarboxylate transporters *SLC16A1* and *SLC16A3* (MCT1 and MCT4^11^) as well as between the mitochondrial iron transporters *SLC25A28* and *SLC25A37* (known as mitoferrin 2 and 1, respectively, Extended Data Fig. 1g). Validation of these interactions using a Multicolor Competition Assays (MCA, Extended Data Fig. 2a-b) confirmed their relevance in HAP1 and A-549 cells, a lung carcinoma cell line, suggesting that these interactions are conserved across different cell lineages (Extended Data Fig. 2c-d). We further validated a second subset of interactions in HAP1 cells, including an interaction between the putative mitochondrial folate/FAD transporter gene *SLC25A32* and *SLC52A2*, a flavin transporter, as well as interactions between the thyroid hormone transporters *SLC16A2* (MCT8) and *SLC16A10* (MCT10) (Extended Data Fig. 2e). Interestingly, thyroid hormones have been previously reported to be required for optimal cancer cell growth^24^, suggesting that these two transporters play functionally redundant roles in HAP1 cells. Moreover, we validated two buffering interactions connecting the cystine/glutamate antiporter *SLC7A11* to the mitochondrial transporters *SLC25A3* and *SLC25A51* (Extended Data Fig. 2f).

**Fig. 2.**
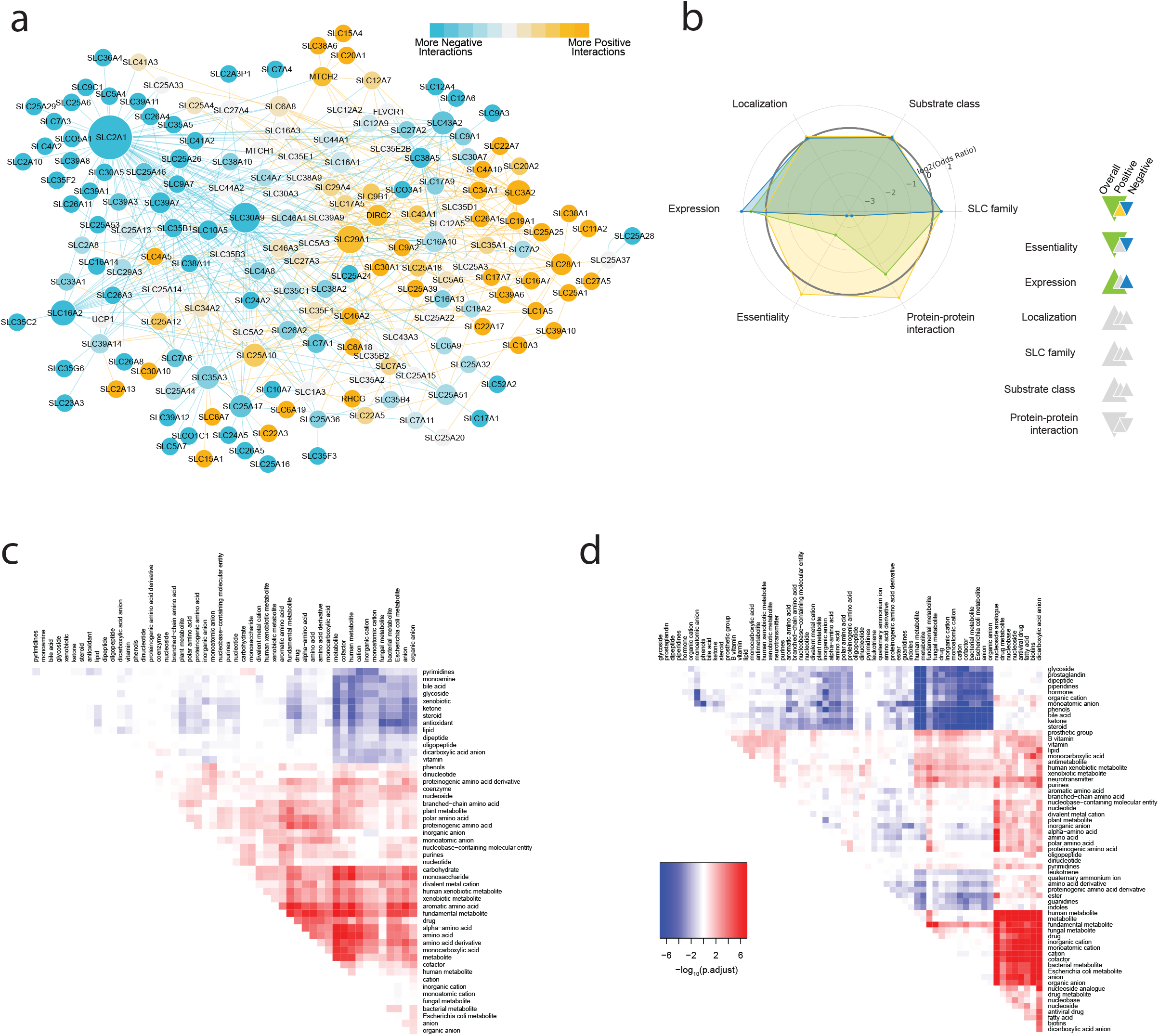
Characterization of a set of strong SLC-SLC genetic interactions. (a) Network representing the set of SLC-SLC genetic interactions with absolute GIscore value above 7 at T_late_. Negative interactions are shown as blue edges, positive as yellow edges. Node size reflects the degree of connectivity. (b) Enrichment analysis with the set of SLC-SLC interactions shown in (a) for Localization, Expression, Essentiality, Protein-Protein Interactions (PPIs), Substrate class. The larger triangles reflect positive (upward) or negative (downward) enrichments for the whole set, the smaller triangles for the positive (yellow) or negative (blue) interactions subsets. For interactions with p<0.05 the corresponding colors are shown. (c, d) Heatmaps showing enrichment (red) and depletion (blue) of interactions between substrate ontology classes in the subset of negative (c) and positive (d) strong interactions.

In order to identify potentially informative patterns within our dataset, we employed a careful annotation of all the members of the human SLC family derived from the primary literature (Extended Data Fig. 4a-b)^25^. As expected, we observed an enrichment for genes expressed in HAP1 within the subset of strong interactions (Fig. 2b, Extended Data Fig. 3b). No overall enrichment was observed for categories such as localization, subfamily or substrate class. However, closer inspection of the subset of strong negative interactions revealed an enrichment of interactions involving the SLC25 family of mitochondrial transporters with the SLC2 (carbohydrate) and SLC43 (amino acids and nucleobases) families (Extended Data Fig. 4d), possibly reflecting their intertwined roles in cellular energetics. Among strong positive interactions, we observed an enrichment for interactions involving nucleoside/nucleotide transporters including members of the SLC28, SLC29, SLC35, as well as the SLC49 family of heme transporters (Extended Data Fig. 4e). We also detected an enrichment of interactions among transporters localizing to the ER and Golgi within the strong negative interactions (Extended Data Fig. 4f-g). To perform finer-level analyses with respect to the substrates of genetically strong interacting transporters, we applied a substrate-based ontology obtained by mapping annotated SLC substrates to the ChEBI ontology (Extended Data Fig. 4c)^25^. This unveiled an enrichment for negative interactions among amino acid transporters, suggesting functional redundancy within this subclass, as well as an enrichment for positive interactions between nucleobase/nucleotide transporters and anion/cation transporters (Fig. 2c-d), therefore providing a rich set of functional connections for future in-depth analyses and validation studies.

Correlation of genetic interaction profiles has been successfully used to define functionally related genes in large genetic interaction datasets^6,8^. We reasoned that exploration of the genetic interaction profile of any given gene, as well as of direct gene-gene interactions, including the annotated functional information, should allow us to infer novel functional annotations (Fig. 3a). To extend our search for interesting patterns and potential annotation of orphan SLCs, we applied a network-based enrichment approach to our transporter dataset and generated a network of profile similarities among SLC genes. Network-based reduction of the total network to retain the most significant connections^26^ highlighted several distinct neighborhoods of SLCs enriched in specific substrates and substrate classes, subcellular localizations and metabolic pathways (Fig. 3b, Extended Data Fig. 5b). Importantly, by employing the functional annotation described above and a neighborhood-based enrichment algorithm (Extended Data Fig. 5a), we could annotate orphan SLCs by analyzing their context in network structure based on their local neighbors using genetic interaction profile similarities (Fig. 3c, Extended Data 5c) or direct gene-gene interactions (Fig. 3d) networks. Several SLCs showed neighborhood enrichments consistent with their family or the functions described in orthologous proteins. These include the orphan gene *SLC38A10*, which connects the two major regions of the network (Fig. 3e), and whose neighborhood is enriched for transporters of proteogenic amino acids (p = 0.0026), consistent with the ability to transport glutamine, glutamate and aspartate reported for its murine orthologous protein^27^. Similarly, the *SLC12A9* neighborhood was enriched for cation-coupled chloride transporters and K+ transport terms (Fig. 3f, p = 0.00012). The SLC12 family is composed of electroneutral cation-coupled chloride transporters, with the subset of SLC12A4-7 being potassium-coupled (KCCs)^28^. Our data suggests therefore that SLC12A9 is an additional member of the KCCs subfamily, possibly localized at the lysosome due to its position within the overall network (Fig. 3b). A similar functional annotation (Reactome R-HSA-426117, p = 2.4 10^−5^) could also be ascribed to the orphan *SLC35E2B* (Fig. 3g).

**Fig. 3.**
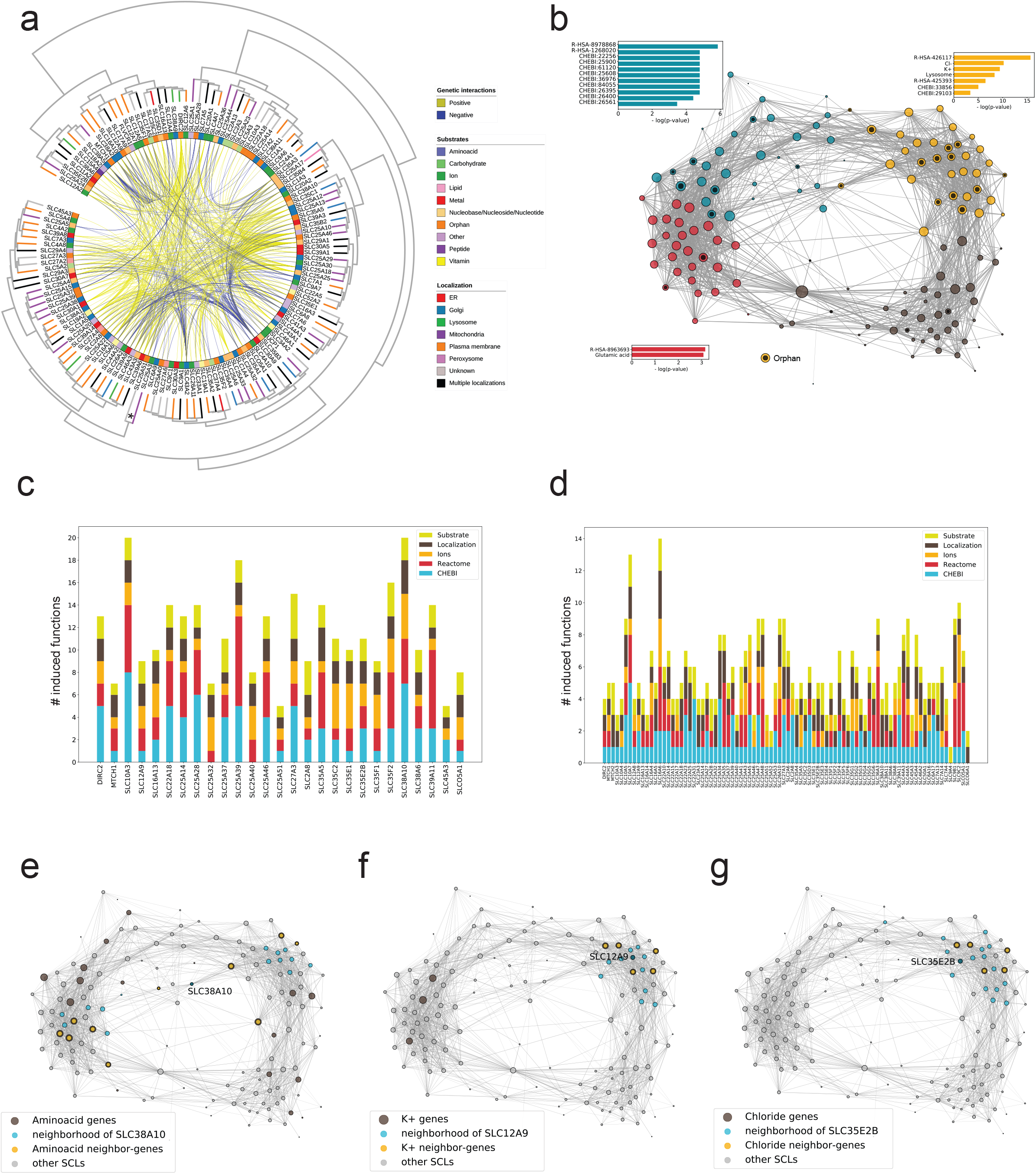
Network-based annotation of orphan SLCs. (a) Integration of the genetic interaction data with substrate and localization annotations. Genes present in the SLC KO collections are arranged on a circle and clustered by genetic interaction profile similarity. Direct gene-gene interactions are shown as connections within the circle. Substrate classes are annotated in the inner color band. Localization is shown on the dendrogram leaves. The * symbol denotes the position of the cluster composed by the mitochondrial *SLC25A3* and *SLC25A51* genes. (b) Network showing the SLCs in the HAP1 KO collection linked by genetic interaction profile similarity. Annotations enriched in the corresponding neighborhoods are shown with the corresponding p value. (c) Plot showing the type and number of inferred functions associated to each orphan SLC in the HAP1 SLC KO collection based on the profile similarity network. (d) Same as in C but based on strong gene-gene interactions. (e-g) Example of orphans SLCs and their neighborhoods, including SLC38A10 (e), SLC12A9 (f), SLC35E2B (g).

Within the dataset, the strongest measured interactions involved a small network of transporters centered around the orphan gene *SLC25A51*, encoding Mitochondrial Carrier Triple Repeat 1 (MCART1, Fig. 4a). Network-based annotation of SLC25A51 suggested nucleosides/nucleotides and phosphate as potential substrates (Extended Data Fig. 5d) as well as a mitochondrial localization (Extended Data Fig. 5e). Interestingly, we observed a similar set of genetic interactions for *SLC25A51* and *SLC25A3*, a phosphate/copper mitochondrial transporter which has been reported to be required for the biogenesis of complex IV of the electron transport chain^29^, consistent with their similar genetic interaction profiles across the full dataset (Fig. 3a). Both genes showed negative interactions with *SLC2A1*/GLUT1, the major glucose transporter at the plasma membrane, as well as positive interactions with *SLC7A11* and *SLC3A2*, the two genes whose corresponding proteins form the heterodimeric transporter xCT, a glutamate/cystine transporter also expressed at the plasma membrane^30^. We confirmed these positive interactions experimentally in the case of *SLC7A11* (Extended Data Fig. 2f). Prompted by this profile similarity, we tested whether *SLC25A51*-deficient cells showed a defect in mitochondrial respiration. When measuring the Oxygen Consumption Rate (OCR), *SLC25A51*-deficient cells showed a loss of mitochondrial respiration comparable to *SLC25A3*-deficient cells (Fig. 4b, Extended Data Fig. 6a). Importantly, ectopic expression of *SLC25A51*, but not *SLC25A3*, restored respiration to levels similar to wt cells, suggesting that the two transporters affected OCR in non-redundant ways, likely through the transport of different substrates (Fig. 4b, Extended Data Fig. 6b). Consistent with the OCR results, a co-dependency analysis^31^ based on the profile of essentiality of *SLC25A51* across the DepMap dataset^32^ revealed *SLC25A3* as the most correlated SLC, as well as high correlation to genes involved in the electron transport chain, ATP synthase and mitochondrial ribosome (Extended Data Fig. 6c-d). To gain further insights into the metabolic effects of the loss of *SLC25A51* and *SLC25A3*, we characterized the metabolic changes in cells lacking one of these two genes compared to wt cells by measuring the abundances of a panel of 194 metabolites by LC-MS/MS (Fig. 4c, Extended Data 6f). We observed a similar pattern of changes (Extended Data Fig. 6g), including depletion of riboflavin, purine nucleotides and enrichment of AICAR, a precursor of IMP, as well as increased levels of metabolites associated to glutathione metabolism, pointing to a defect in one carbon metabolism (Fig. 4d). This is consistent with the observed negative interactions of *SLC25A51* with the putative mitochondrial folate/FAD transporter gene *SLC25A32* as well as the purine transporter gene *SLC43A3*, which has been previously linked to the purine salvage pathway (Fig. 4a)^33^. Interestingly, comparison of changes in metabolites between SLC25A51-and SLC25A3-deficient cells showed stronger depletion of TCA cycle intermediates, including citrate, aconitate and cis-aconitate, isocitrate and succinate, in SLC25A51 knockouts (Fig. 4e, Extended Data Fig. 6g). Moreover, analysis of the co-dependencies specific to *SLC25A51* and not *SLC25A3* highlighted a similarity to mutations of pyruvate dehydrogenase and citrate synthase (Extended Data Fig. 6e), suggesting that loss of SLC25A51 mimicked the loss of enzymes acting at the early stages of the TCA cycle. We therefore reasoned that this transporter could provide a small, nucleoside-containing (Extended Data Fig. 5e) molecule or cofactor involved with both the early stages of the TCA cycle as well as the assembly/functioning of the ETC (Fig. 4e). Among these cofactors and vitamins, nicotinamide adenosine dinucleotide (NAD) was the only significantly depleted molecule in SLC25A51-deficient cells versus SLC25A3-deficient ones (Fig. 4c, Extended Data 6f-g), leading us to hypothesize that SLC25A51 could be involved in the uptake of NAD+ or its precursors. Consistent with these results, NAD(H) levels were reduced in SLC25A51-deficient cells compared to SLC25A3 knockouts, when measured by a luminescence-based assay (Extended Data Fig. 6h). It has been recently shown that mitochondria can import NAD(H)^34^ and that the cytosolic NAD(H) levels affect the mitochondrial pool^35^, suggesting the presence of a transporter responsible for NAD compartmentalization^36^. We reasoned that, if *SLC25A51* encodes a NAD(H) transporter function in human cells, the well characterized *Saccharomyces cerevisiae* mitochondrial NAD+ transporters, Ndt1p and Ndt2p^37^, should be able to functionally rescue the defect. Indeed, while no expression was observed for *NDT2* (Extended Data Fig. 6i), ectopic expression of yeast *NDT1* restored the mitochondrial respiration defect of the SLC25A51-deficient cells (Fig. 4f, Extended Data Fig. 6i). While additional data, including measurement of mitochondrial metabolites levels and transport assays with reconstituted protein, will be required to definitely prove the role of SLC25A51 as NAD+ transporter, the multiple functional and metabolic links described above support a role for SLC25A51 in mitochondrial NAD(H) metabolism and highlight the potential of using genetic interactions to functionally annotate orphan genes.

**Fig. 4.**
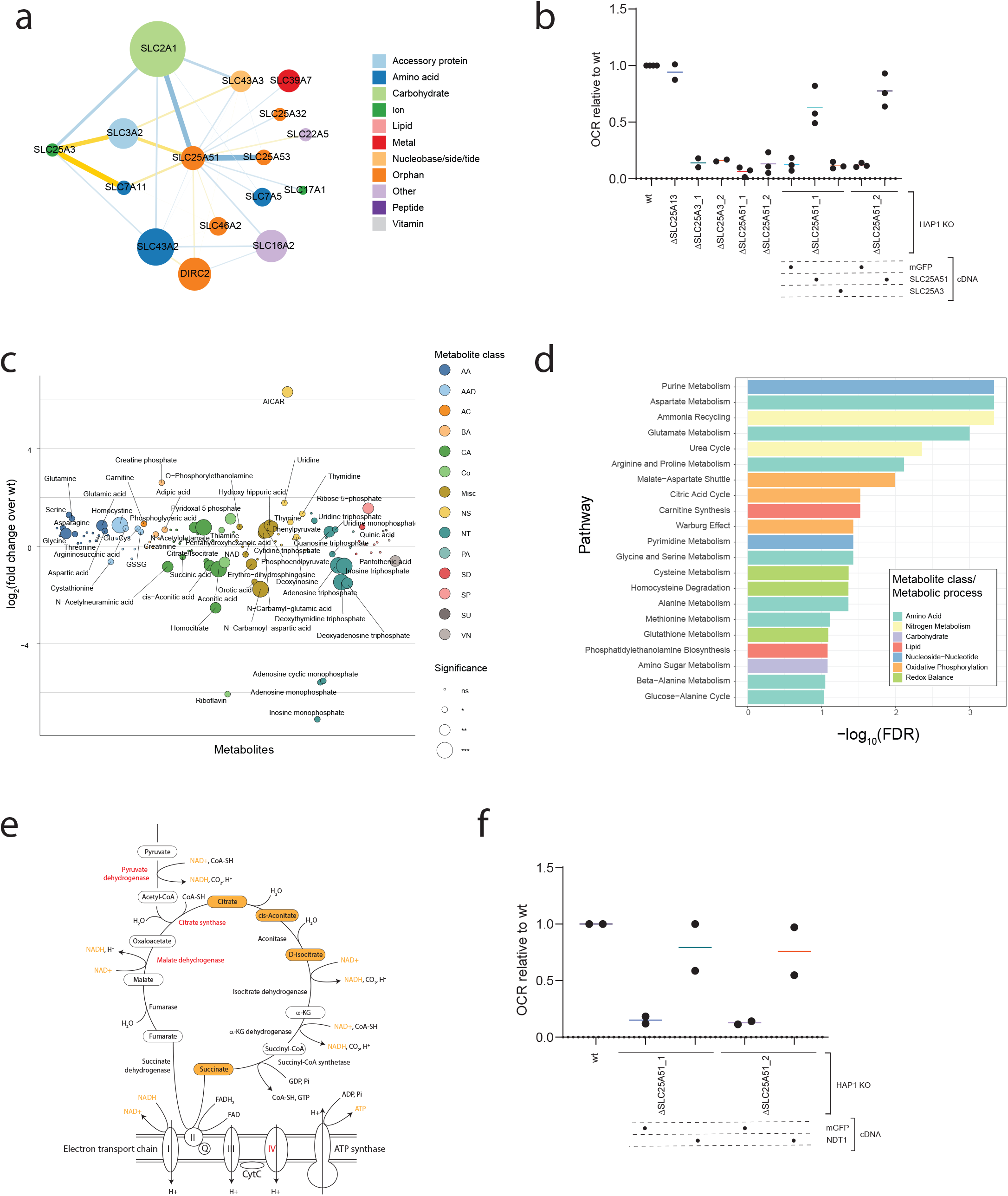
Functional deorphanization of SLC25A51 reveals a role in mitochondrial NAD(H) transport. (a) Sub-network of strong genetic interactions involving the orphan gene SLC25A51. Node color refers to substrate class, node size reflects the degree of connectivity in the strong SLC-SLC gene interaction network. (b) Oxygen consumption rate (OCR) in wt and SLC-deficient HAP1 cells, as well as cells reconstituted with the indicated cDNAs. mGFP: mitochondrially-localized GFP. SLC25A13 KO cells are used as negative control. Results of 2-3 independent biological replicates are shown. The bars show average value across biological replicates. (c) Targeted metabolomics profile of SLC25A51-deficient cells compared to wt HAP1 cells. Metabolite classes are indicated by different colors (AA – Amino Acids, AAD – Amino Acid Derivatives, AC – Acylcarnitines, BA – Biogenic Amines, CA – Carboxylic acid, Co – Cofactors, Misc – miscellaneous, NS – Nucleosides/Nucleobases, NT – Nucleotides, PA – Phenyl Acids, SD – Sugar derivates, SP – Sugar phosphate, SU – Sugars, VN – Vitamin). Circle sizes reflect significance of the log_2_ fold change measured (*** pvalue <0.01, ** pvalue <0.05, * pvalue <0.1, ns non-significant). (d) Enrichment analysis of metabolic pathways affected in SLC25A51 KO cells compared to wt cells, using the SMPBD database as reference. (e) Schematic view of the TCA and ETC pathway/complexes in the mitochondria. Metabolites depleted in SLC25A51 but not in SLC25A3 KOs are shown in orange boxes. Enzymes with similar essentiality profiles as SLC25A51 across the DepMap dataset are shown in red. (f) Oxygen consumption rate (OCR) in wt and SLC25A51-deficient HAP1 cells reconstituted with mitochondrially-localized GFP (mGFP) or the yeast NAD+ transporter *NTD1*. Results of 2 independent biological replicates are shown. The bars show average value across biological replicates.

Despite the high degree of functional redundancy displayed by transporters^38^, we show here that informative patterns of genetic interactions can be systematically derived for this family, allowing the generation of testable hypotheses for the de-orphanization of poorly characterized SLCs. Further studies employing higher-order genetics^10,11,39^ or modifications of media composition^40^ can be expected to reveal further dependencies among this class of genes. As the functional breadth of SLCs and their potential as a target class becomes increasingly recognized^1,4^, systematic approaches such as the genetic interaction map described here will be necessary to define functions and therapeutically relevant applications of Solute Carriers. Mapping genetic interactions in human cells promises to be a very powerful approach to assign gene products to core biochemical and cellular functions. While a high accuracy genome-wide screen may still be technically challenging and an annotation purgatory, a dissection function-by-function, as proposed here for transmembrane transport, bears the advantage to foster a comparative and integrative dimension allowing abundant assignment of functions through guilt-by-association as well as exclusion criteria. The study presented here may thus act as a blueprint for the systematic deorphanization of genes involved in related cellular processes.

---

**Extended Data Fig. 1.**
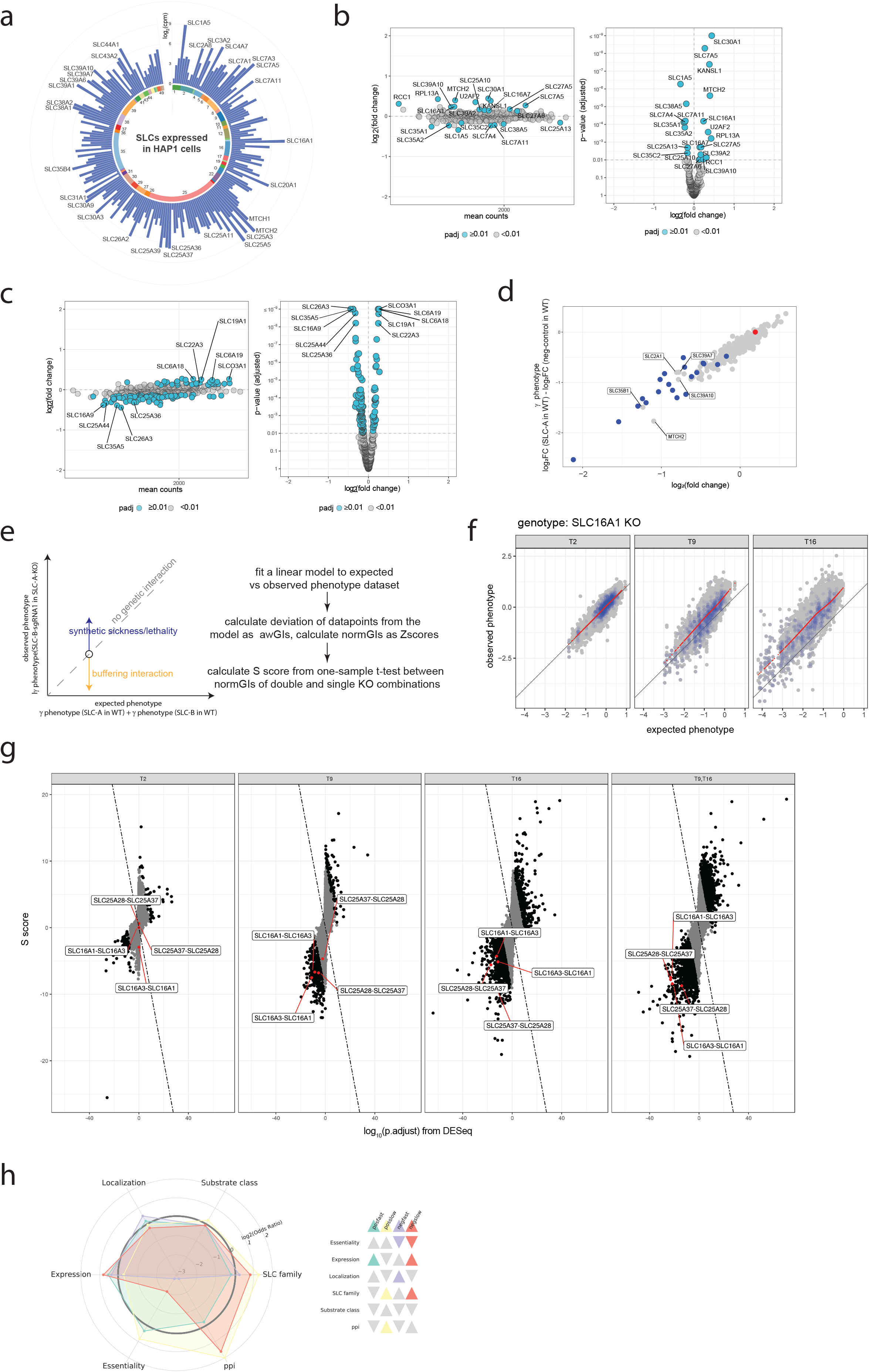
Genetic interaction scoring. (a) Expression level of the SLCs targeted by the CRISPR/Cas9 KO library in HAP1 cells. Genes are arranged by subfamily, indicated by the number in the circle. Bars show the expression of the corresponding transcripts in HAP1 cells (count per million reads, cpm). Genes with expression above 2^6^ cpm are labeled. (b) Magnitude-amplitude (left) and volcano (right) plots of the genetic interaction screen with *DBT*-deficient cells at 16 days p.i. Mean counts per gene targeted are shown as well as log2(fold changes) over the wt samples. Adjusted pvalues were calculated with DESeq. Significant genes (adjusted p value < 0.01) are shown as blue circles. (c) Magnitude-amplitude (left) and volcano (right) plots of the genetic interaction screen with *CPS1*-deficient cells at 16 days p.i. Mean counts per gene targeted are shown as well as log2(fold changes) over the wt samples. Adjusted pvalues were calculated with DESeq. Significant genes (adjusted p value < 0.01) are shown as blue circles. (d) Plot of log2FC and γ phenotype at the gene-level (median of sgRNA-level effects) for wt cells at 16 days p.i. Positive control essential genes are shown as blue dots, the negative control as a red dot. (e) Scheme of the S score calculation. (f) Plot of expected versus observed γ phenotypes for all sgRNAs at the indicated timepoints for the screen against the SLC16A1-deficient genotype. sgRNAs targeting positive control genes are shown as blue dots. The black line indicates the equivalence of expected and observed phenotypes. The red dots indicate the model predictions in bins of 200 expected phenotypes each. (g) Plot comparing the adjusted p values obtained from DESeq2 and the S scores at the gene level. The line corresponds to interactions for which the average of the two score is 0 (GIscore = 0). Interactions with |GIscore| >5 are shown in black. Two sets of reciprocal interactions involving SLC16A1-SLC16A3 and SLC25A28-SLC25A37 are shown in red. (h) Enrichment analysis with the set of fast and slow genetic interactions for Localization, Expression, Essentiality, PPIs, Substrate class, SLC family. The triangles reflect positive (upward) or negative (downward) enrichments for the whole set or the specific category listed.

**Extended Data Fig. 2.**
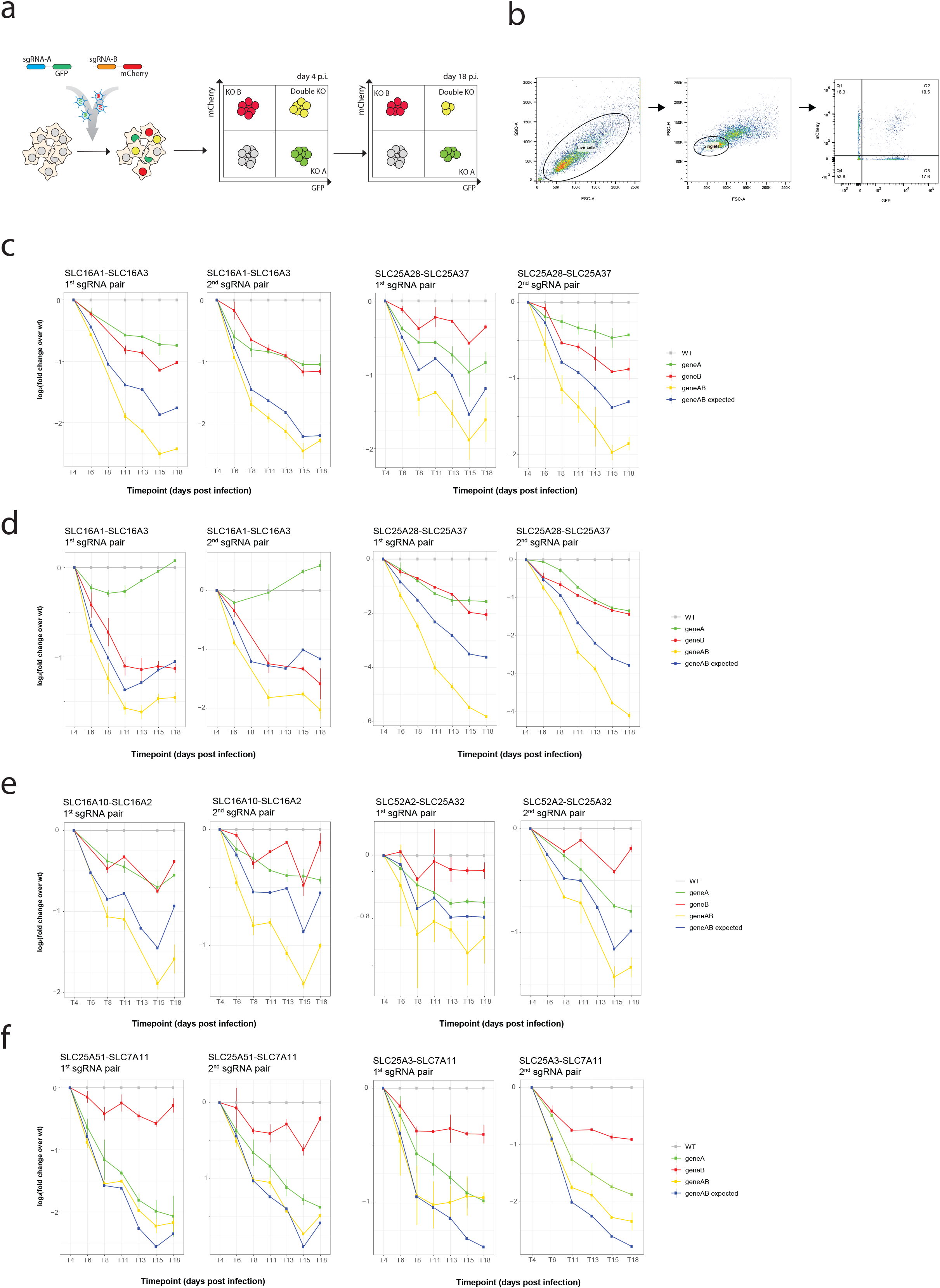
Validation of SLC genetic interactions by Multicolor Competition Assay (MCA). (a) Scheme of the experimental setup for the MCA. (b) Example of gating strategy for a MCA experiment. (c)(e)(f) Log_2_(fold change) of population with single or double KOs compared to wt in MCA assays in HAP1 cells. Two combinations of sgRNA for each gene pair were used. Points indicate the average of two technical replicates. Vertical lines indicate the values for each technical replicate. One representative experiment out of two independent replicates is shown. (d) Same as (c) but in A-549 cells.

**Extended Data Fig. 3.**
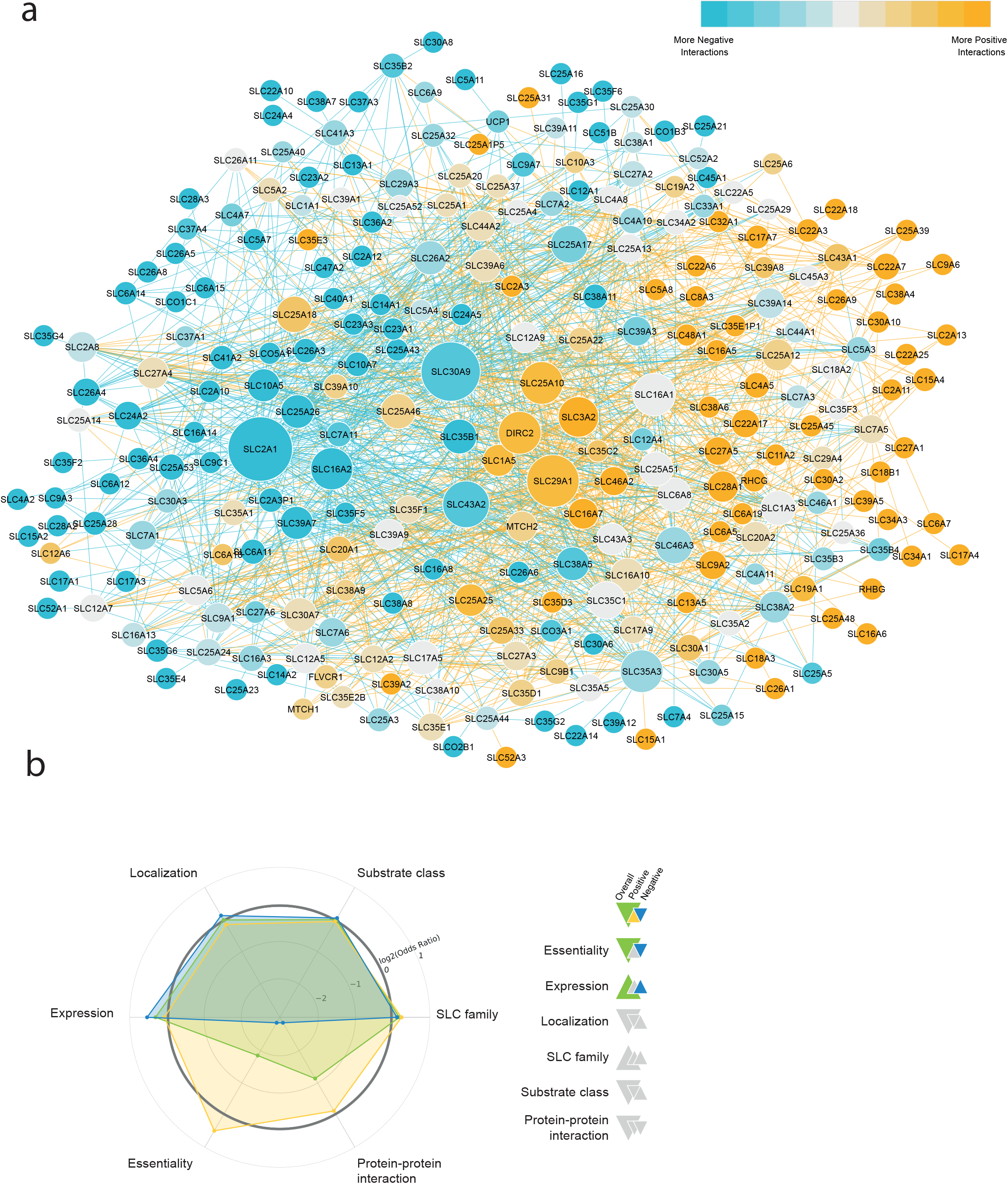
Network of strong genetic interactions. (a) Network representing the set of SLC-SLC genetic interactions with absolute GIscore value above 5 at T_late_. Negative interactions are shown as blue edges, positive as yellow edges. Node size reflects the degree of connectivity. (b) Enrichment analysis with the set of SLC-SLC interactions shown in (a) for Localization, Expression, Essentiality, Protein-Protein Interactions (PPIs), Substrate class. The larger triangles reflect positive (upward) or negative (downward) enrichments for the whole set, the smaller triangles for the positive (yellow) or negative (blue) interactions subsets. For interactions with p < 0.05 the corresponding colors are shown.

**Extended Data Fig. 4.**
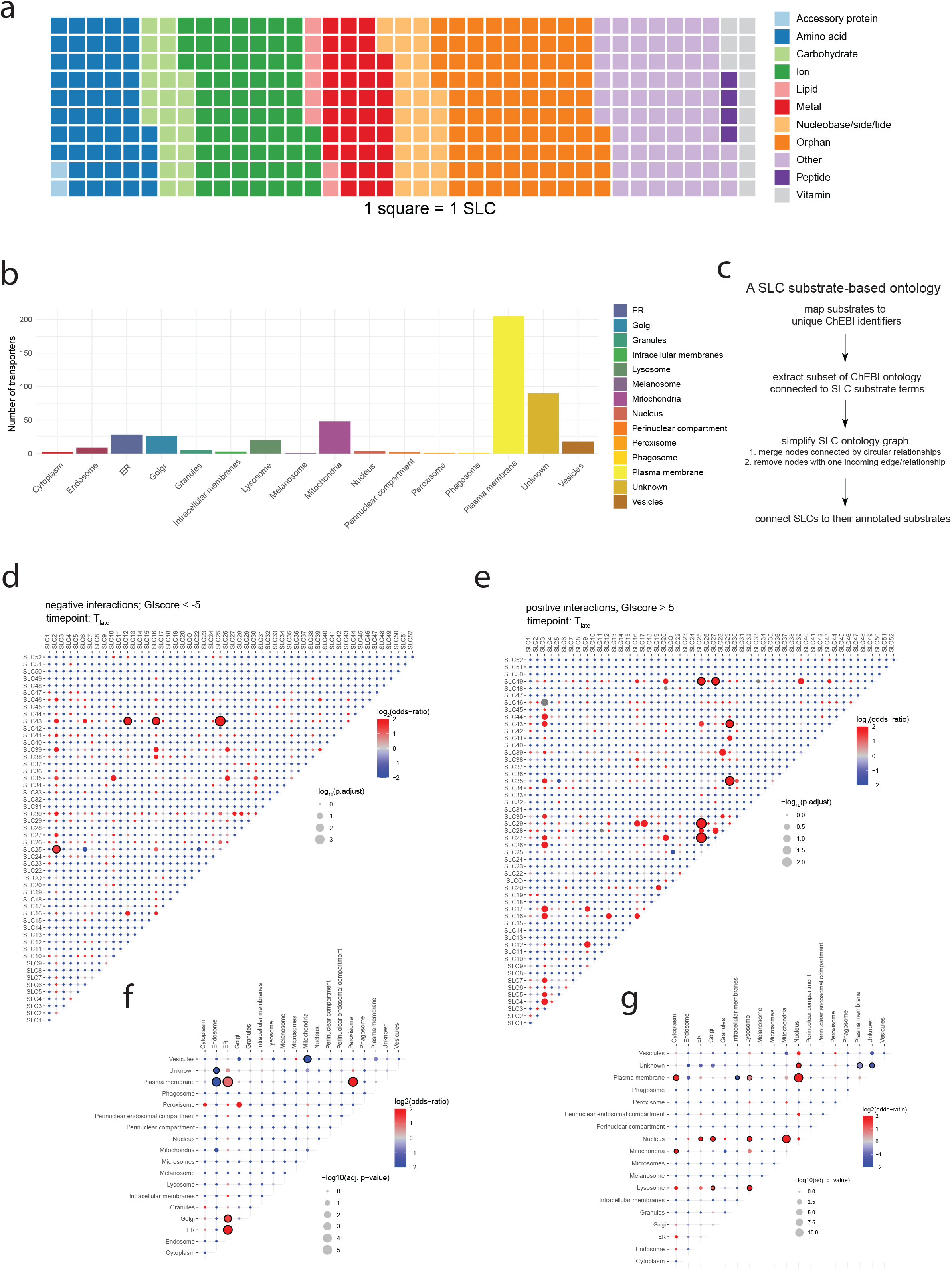
SLC annotation and annotation enrichments. (a) Substrate class annotation for the SLCs present in the SLC KO CRISPR/Cas9 library. (b) Localization annotation for the SLCs present in the SLC KO CRISPR/Cas9 library. (c) Scheme of the substrate-based ontology generation. (d) Enrichment at the family-level within the subset of interactions with GIscore < −5. (e) Same as in (d) for interactions with GIscore > 5. (f) Localization-level enrichments within the subset of interactions with GIscore < −5. (g) Same as in (f) for interactions with GIscore > 5.

**Extended Data Fig. 5.**
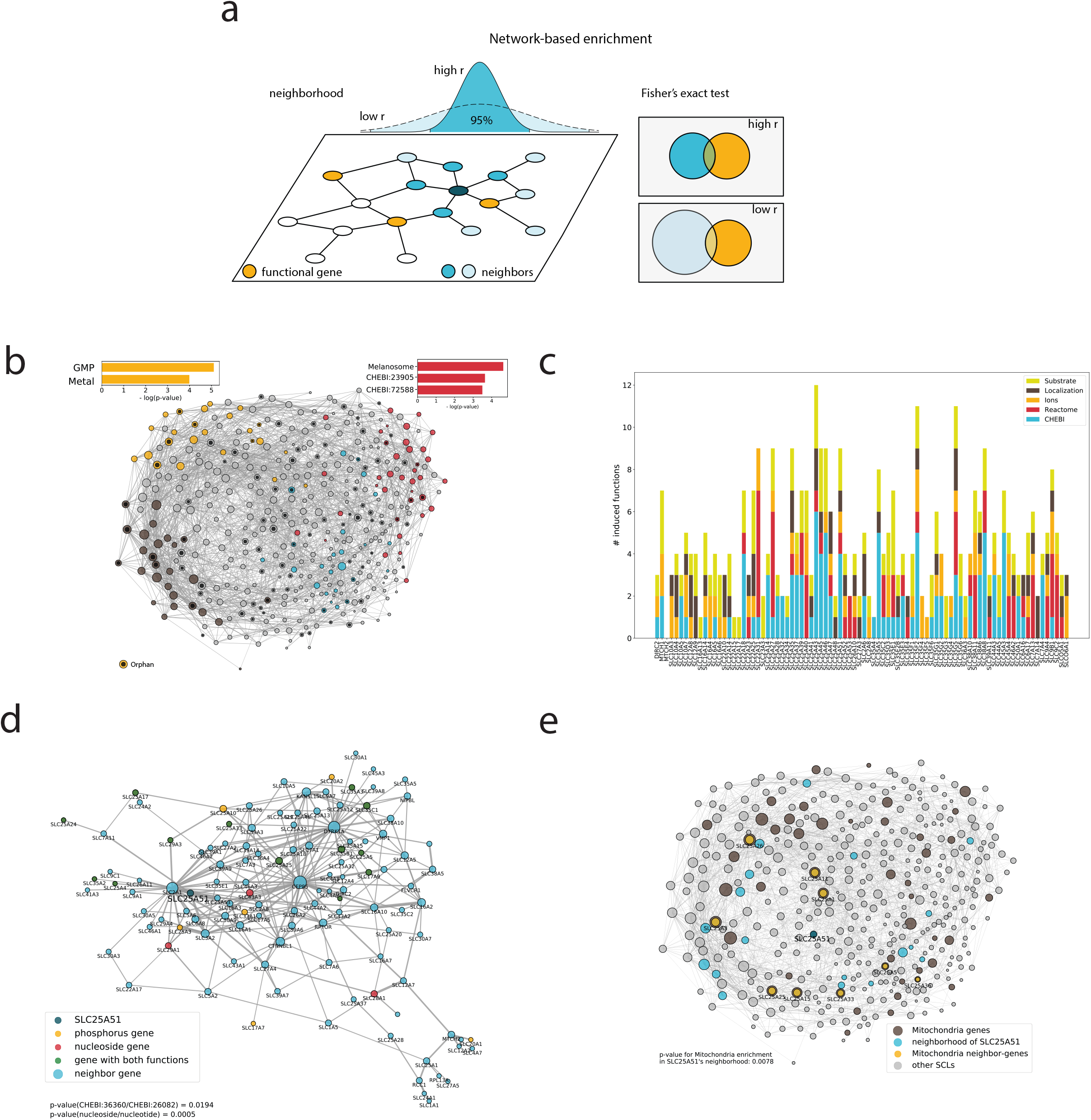
Genetic interaction profile similarity SLC networks (a) Network-based enrichment: A random walk with restart initiated from an orphan gene assigns visiting frequencies for all genes in the network. The number of neighbor genes is given by the set of nodes containing 95% of the overall visiting frequency. A restart value r close to one results in a small neighborhood (blue nodes), densely localized around the orphan, whereas low values of r generate a broader range of neighbors (light blue nodes). This set is tested for enrichment of a specific function (yellow nodes). Significantly enriched functions in the neighborhood are considered to be induced orphan functions. (b) Network showing SLCs targeted in the CRISPR/Cas9 library, linked by genetic interaction profile similarity. Functions enriched in the corresponding neighborhoods are shown with the corresponding p value. (c) Plot showing the type and number of induced functions associated to each orphan SLCs, derived from the network shown in (B). (d) Neighborhood of the orphan gene SLC25A51, derived from the row-based network shown in Fig. 3B. (e) Neighborhood of the orphan gene SLC25A51, derived from the column-based network shown in (b).

**Extended Data Fig. 6.**
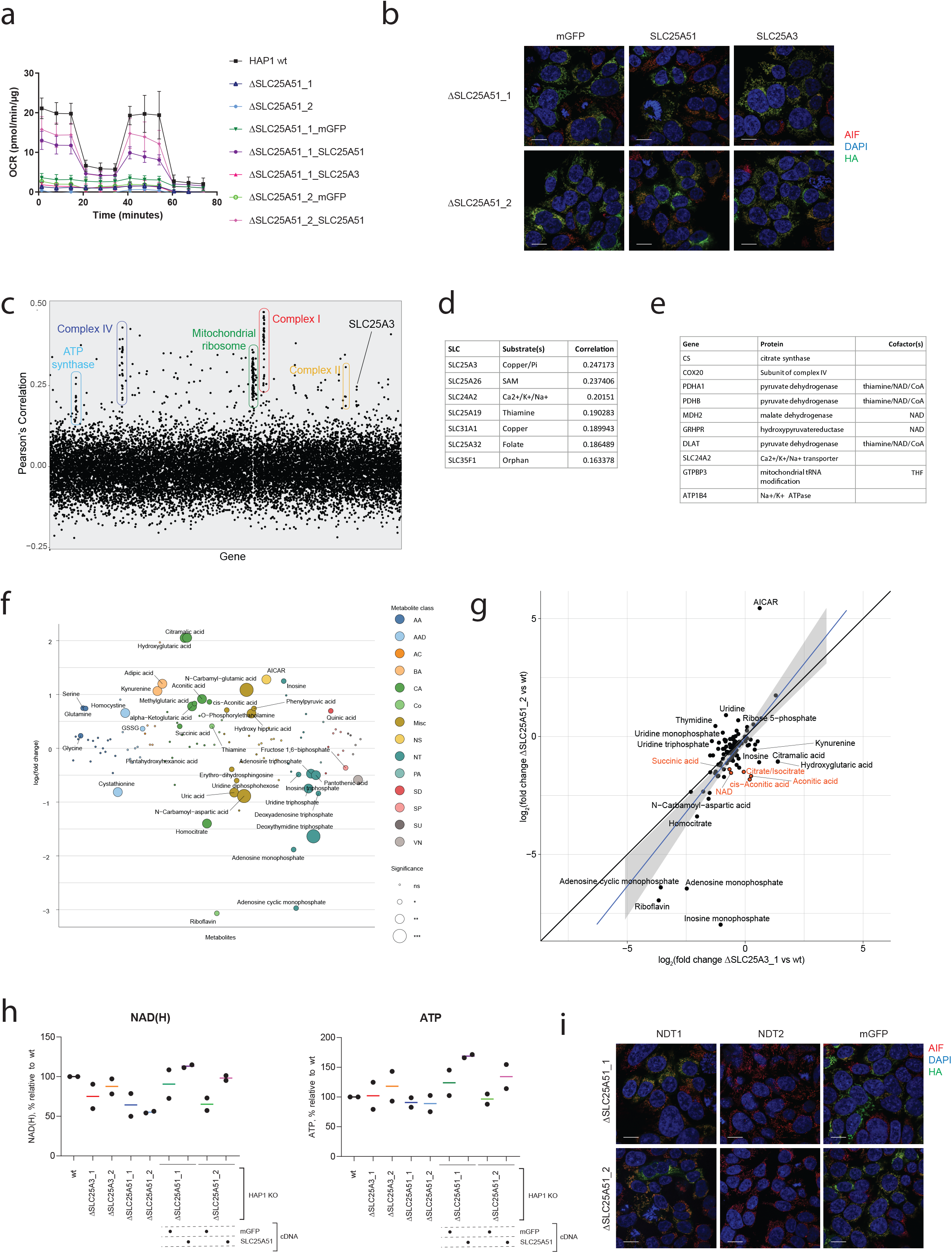
SLC25A51 as mitochondrial NAD transporter. (a) Oxygen consumption rate from a representative Seahorse measurement. Average and standard deviation of at least 6-8 technical replicates. (b) Representative confocal images of HAP1 cells lacking endogenous SLC25A51 and reconstituted with mitoGFP, SLC25A3 or SLC25A51 cDNA. Green: ectopic HA-tagged protein, Blue: DAPI, Red: Apoptosis inducing factor (AIF). Scale bar: 20μm. (c) Plot of the correlations of essentiality across cell lines in the DepMap dataset for each human gene, in relation to SLC25A51. Members of major mitochondrial complexes, as well as the SLC with the most highly correlated essentiality profiles are labeled. (d) List of the SLCs with the most highly correlated essentiality profiles in relation to SLC25A51. (e) List of the top correlated genes specific to SLC25A51 but not SLC25A3. (f) Targeted metabolomics profile of SLC25A3-deficient cells compared to wt HAP1 cells. Metabolite classes are indicated by different colors (AA – Amino Acids, AAD – Amino Acid Derivatives, AC– Acylcarnitines, BA – Biogenic Amines, CA – Carboxylic acid, Co– Cofactors, Misc– miscellaneous, NS – Nucleosides/Nucleobases, NT– Nucleotides, PA – Phenyl Acids, SD – Sugar derivates, SP – Sugar phosphate, SU – Sugars, VN – Vitamin). Circle size reflect significance of the log_2_(fold change) measured (p adjusted * < 0.05, ** < 0.01, ***< 0.001, ns non-significant). (g) Comparison of the log_2_(fold change) of metabolite amounts in SLC25A51- and SLC25A3-deficient cells compared to wt. Diagonal line y = x is shown in black, linear fit to the data is shown in blue with grey shaded area corresponding to 95% confidence interval. Metabolites differentially affected in the two Kos are labeled, with the subset involved with TCA cycles labeled in red. (h) Total cellular levels of NAD/NADH (lefthand panel) and ATP (righthand panel) in the indicated cell lines, normalized to wt levels. Cumulative results of two biological replicates. (i) Representative confocal images of HAP1 cells lacking endogenous SLC25A51 and reconstituted with HA-tagged *mitoGFP, NDT1* or *NDT2* cDNA. Green: ectopic HA-tagged protein, Blue: DAPI, Red: AIF. Scale bar: 20μm.

## Materials and Methods

### Cell lines and reagents

HAP1 wt cells as well as single cell-derived clones were obtained from Haplogen Genomics or generated in-house by transient transfection with px459 (Addgene #48139) vectors carrying sgRNAs against the selected genes. Two independent clones for each gene were used whenever available. HAP1 cells (Haplogen Genomics) were grown in IMDM with 10% FBS, 1% Pen/Strep. HEK293T and A-549 cells (ATCC) were grown in DMEM with 10% FBS, 1% Pen/Strep (Gibco). For CRISPR-based knockout cell lines, sgRNAs were designed using CHOPCHOP^41^ and cloned into pLentiCRISPRv2 (Addgene, #52961), LGPIG (pLentiGuide-PuroR-IRES-GFP) or LGPIC (pLentiGuide-PuroR-IRES-mCherry)^42^. sgRen, targeting the *Renilla* spp. luciferase gene, was used as negative control sgRNA (38). The SLC-deficient clones (ΔSLC25A51_1927-10, ΔSLC25A51_1927-11, ΔSLC25A3_792_1, ΔSLC25A3_792_6, ΔSLC25A13_789_6, renamed as ΔSLC25A51_1, ΔSLC25A51_2, ΔSLC25A3_1, ΔSLC25A3_2, ΔSLC25A13_1) were obtained from Haplogen Genomics. Codon-optimized SLC25A51, SLC25A3 cDNAs or mitoGFP cDNA sequences (the latter carrying the mitochondrial import sequence derived from subunit 8 of complex IV) were obtained from the ReSOLUTE consortium (https://re-solute.eu/). The yeast NAD+ transporters ndt1/yil006w and ndt2/yel006w were obtained from Horizon Discovery (catalog IDs YSC3867-202327426 and YSC3867-202326304, respectively). cDNAs were cloned into a modified pCW57.1 lentiviral vector (Addgene #41393) generated within the ReSOLUTE consortium and carrying a Strep-HA tag and blasticidin resistance.

### Genetic screening

Genetic screening was performed by transducing wild type or knockout cell lines with the SLC CRISPR/Cas9 library^15^ at an MOI of 0.2-0.3 and 2000x coverage, in triplicate, followed by selection with puromycin (1μg/ml) for 7 days. Samples were collected at 2, 9 and 16 days post-infection, the genomic DNA extracted with QIAGEN DNA Easy kit. Amplification was performed by PCR using Q5 polymerase (NEB) for 26-28 cycles using primers carrying dual indexes as previously described^15^. Samples were normalized based on band intensity and multiplexed together to achieve a coverage of approximately 2000x by NGS on an Illumina 3000/4000 platform (BSF facility, CeMM/MUW).

### Sample sequencing and genetic interaction analysis

Reads were demultiplexed and mapped to the sgRNAs present in the library using a custom procedure implemented in Python (crispyCount). Samples with a median sgRNA coverage of less than 100 were excluded from further analysis (174 of 2488 samples). Replicates with sgRNA read counts correlation (Pearson’s) lower than 0.5 to other replicates (224 samples), as well as pair of clones with correlation lower than 0.5 for the same time point (11 samples) were also excluded from further analysis. Six wt samples (out of a total of 54 wt sample) showing outlier distributions upon manual inspection were further excluded. In total, 415 of 2103 samples were excluded. An additional time point (T_late_) was created by merging the data points from the T_9_ and T_16_ time points. A set of 100 negative sgRNAs was selected (from the initial group of 120 negative control sgRNAs) based on the lack of fitness phenotype in a previous genetic screen performed in HAP1 cells with the same library^15^. Statistical enrichment or depletion of genes in a specific genetic background as compared to wild type was calculated in R, employing the DESeq2 library with sgRNA read count tables as input for creating a linear model per gene with factors for individual guides, time points, and clones. Resulting p-values were corrected for multiple testing using the Benjamini-Hochberg method^43^. Custom normalization factors were calculated, assuring equal median sgRNA read count per sample. In parallel, a S score was calculated as described in ^44^. Briefly, an enrichment factor was calculated as the log_2_ of the ratio of normalized counts for each sgRNA at a given time point compared to the original plasmid library. γ phenotypes were then calculated by subtracting from the enrichments of sgRNAs targeting genes the median enrichment of negative control sgRNAs, for each clone/timepoint/replicate combination. Expected phenotypes were calculated as the sum of the γ phenotypes derived for each gene in the wild type samples. Observed and expected phenotypes were plotted against each other and a generalized additive model (GAM) was used to define the typical behaviour of the measured gene pair combinations at each time point. Raw genetic interactions (GI) were calculated as the residuals between the measured γ phenotype and the model prediction, and further normalized by dividing them by the standard deviation of the Raw-GIs of the neighboring 200 observations along the expected phenotype axis to yield normGIs. A S score, equivalent to a Student’s t score, was then calculated as follows:

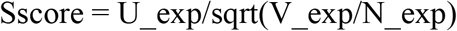

Where:

U_exp = median of normGIs for each gene pair

N_exp = number of sgRNA combinations targeting each gene pair

V_exp = variance of normGIs for each gene pair

The adjusted p-values from DESeq2 and the S scores were finally linearly combined to generate a single genetic interaction score using the formula:

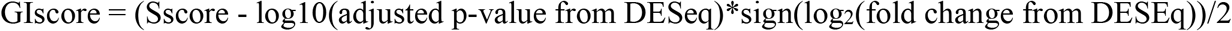

### Genetic interaction time course

Genetic interaction time courses were plotted using the GIscores from T2, T9, and T16. To distinguish between fast and slow interactors, the deviation of T9 from the line interpolated between T2 and T16 was calculated:

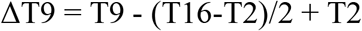

For this calculation only, scores near zero were denoised (GIscore < 1 were set to 0). Additionally, interaction time courses which were not strictly increasing or decreasing were filtered out, as well as interaction pairs that had mixed positive and negative interaction scores. The 250 interaction pairs with the largest ΔT9 are defined as the “fast interactor” group, where the bulk of the genetic interaction occurs between T2 and T9. The 250 interaction pairs with the smallest ΔT9 are defined as the “slow interactor” group.

### Multicolor competition assay

HAP1 cells were infected with viral particles containing LGPIC-sgRNA-A (mCherry-positive, targeting gene A) mixed in 1:1 ratio with LGPIG-sgRNA-B (eGFP-positive, targeting gene B) without any antibiotic selection. The mixed cell populations were monitored up to 18 days p.i. and the respective percentage of viable (FSC/SSC) mCherry-positive and eGFP-positive cells at the indicated time points was quantified by flow cytometry. Samples were analyzed on an LSR Fortessa (BD Biosciences) and data analysis was performed using FlowJo software (Tree Star Inc., USA). Individual ratios were normalized to day 4 measurements and then log transformed.

### Enrichment analyses

A comprehensive set of SLC annotations was derived from the primary literature as described in ^25^. Protein-protein interactions are queried from the Human Integrated Protein Protein Interaction Reference (HIPPIE) database v2.2. The original set of interactions integrated from 10 source databases and 11 studies were filtered to only keep protein coding genes resulting in 16,826 nodes and 367,384 edges. 348 SLCs of the screen were present in the PPI network, creating a subnetwork of 348 nodes and 150 edges^45,46^.

Association between annotations (SLC family, Localization, Essentiality, Expression, SLC ontology terms) and a given interaction type (positive, negative, positive fast, positive slow, negative fast, negative slow) was quantified using Fisher’s exact test. Odds-ratios are plotted in the spider plots and enrichment / depletion is indicated by the direction of the triangles. Significant features (p-value < 0.05) are indicated as colored triangles.

### Network construction

The SLC gene interaction network was constructed from the GIscore (see above). Interactions with a |GIscore| > 5 or 7 are represented as an edge in the network. The profile similarity networks were constructed by first calculating the Pearson correlation between the score profiles for all pairs of tested SLCs (control genes were removed). We then applied the network-based disparity filter^26^ on the resulting all-to-all correlation matrix to identify the most significant interactions. All networks can be explored by the reader via our web interface SIGIL (SLC Interactive Genetic Interaction Landscape) at http://sigil.cemm.at/.

### Network-based enrichment

We used a random walk based strategy to explore the network neighborhood around a given orphan SLC gene and identify the functionally most closely related genes^47^. Starting from the gene of interest, the procedure simulates a walker jumping randomly from node to node along the edges of the network. At every step of the iterative procedure, the walker either jumps back to the starting gene (with restart probability r) or selects one of the current network neighbors with a probability proportional to the weight of the respective edge. The procedure eventually converges to stationary frequencies with which the different nodes in the network are visited by the walker. A high visiting frequency indicates that the respective node is in close network proximity of the particular starting node. We define a neighborhood around a given starting node as the set of nodes containing 95% of the overall visiting frequency. This random walk based neighborhood provides a more fine grained measure than using only directly connected neighbors, as it also takes the connectivity structure between neighbors into account. For the functional characterization of orphan genes, we identified significantly enriched annotations among the genes within their random walk neighborhood (Fisher’s exact test with Bonferroni correction for multiple comparisons). Note, that orphan genes tend to be less close to functional genes on the networks than randomly selected genes. As a result, the overlap of a locally defined neighboring gene set (restart value close to one) with the set of genes associated to a specific function may be too small or zero. For such cases, we extended the neighborhood beyond immediate neighbors by decreasing the restart value while keeping neighborhood size reasonable until the overlap is big enough to find enrichment versus the background of all genes in the network. Additionally, the hypergeometric test is designed such that it produces greater p-values for bigger sample sizes that comes with larger neighborhoods. Therefore, small p-values also indicates local neighborhoods and vice versa.

### Mitostress measurements

Cells carrying inducible constructs were treated with 1μg/ml doxycycline for 24h before plating. To limit effects due to different doubling times across the cells lines tested, cells were seeded in 96-well plates at 40.000 cells/well on the same day of the experiment. Before measurement, medium was exchanged for XF Base Medium (Agilent 102353-100) containing glucose (10 mM), sodium pyruvate (1 mM) and L-glutamine (2 mM) and cells were incubated for 1h at 37°C. Measurements were carried out on a Seahorse XF96 (Agilent) with a MitoStress (Agilent, 103015-100) kit, following the manufacturer’s instructions. Oligomycin, FCCP, and a mix of Rotenone and Antimycin A were injected at desired timepoints at a final concentration of 1 μM, 1.5 μM and 0.6 μM, respectively. After measurement, the medium was removed, cells were lysed, and protein amount was determined by a Bradford Assay. Data were normalized to protein amount and analyzed with Seahorse Wave (Agilent) and Prism (Graph Pad).

### Gene expression analysis (RNA-Seq)

HAP1 cells were plated (2×10^6^ cells per condition, in triplicate). Cells were harvested after 24h and RNA was isolated using the QIAGEN RNeasy Mini kit including a DNase I digest step. RNA-Seq libraries were prepared using QuantSeq 3’ mRNA-Seq Library Prep Kit FWD for Illumina (Lexogen) according to the manufacture’s protocol. The libraries were sequenced by the Biomedical Sequencing Facility at CeMM/MUW using the Illumina HiSeq 4000 platform at the 50 bp single-end configuration. Raw sequencing reads were demultiplexed, and after barcode, adaptor and quality trimming with cutadapt (https://cutadapt.readthedocs.io/en/stable/), quality control was performed using FastQC (http://www.bioinformatics.babraham.ac.uk/projects/fastqc/). The remaining reads were mapped to the GRCh38/h38 human genome assembly using genomic short-read RNA-Seq aligner STAR version 2.592. Transcripts were quantified using End Sequence Analysis Toolkit (ESAT)^48^. Differential expression analysis was performed using independent triplicates with DESeq2 (1.24.0) on the basis of read counts.

### Targeted metabolomics

Cells were plated at 0.8-0.9×10^6^ cells/well in 6-well plates in full media. After 24h, the cells were gently washed with room temperature PBS, transferred to ice and 1.5ml of ice-cold 80:20 MeOH:H_2_O solution was added to each well. The cells were scraped and transferred to a pre-cooled Eppendorf tube, snap-freezed in liquid nitrogen and thawed in ice before being centrifuged at 16,000g for 10 minutes at 4°C. Cell extracts were dried downs using a nitrogen evaporator. The dry residue was reconstituted in 50 µL of water. 10 µL of sample extract was mixed with 10 µL of isotopically labelled internal standard mixture in HPLC vial and used for LC-MS/MS analysis. A 1290 Infinity II UHPLC system (Agilent Technologies) coupled with a 6470 triple quadrupole mass spectrometer (Agilent Technologies) was used for the LC-MS/MS analysis. The chromatographic separation for samples was carried out on a ZORBAX RRHD Extend-C18, 2.1 × 150 mm, 1.8 um analytical column (Agilent Technologies). The column was maintained at a temperature of 40°C and 4 µL of sample was injected per run. The mobile phase A was 3% methanol (v/v), 10 mM tributylamine, 15 mM acetic acid in water and mobile phase B was 10 mM tributylamine, 15 mM acetic acid in methanol. The gradient elution with a flow rate 0.25 mL/min was performed for a total time of 24 min. Afterward, a back flushing of the column using a 6port/2-position divert valve was carried out for 8 min using acetonitrile, followed by 8 min of column equilibration with 100% mobile phase A. The triple quadrupole mass spectrometer was operated in an electrospray ionization negative mode, spray voltage 2 kV, gas temperature 150 °C, gas flow 1.3 L/min, nebulizer 45 psi, sheath gas temperature 325 °C, sheath gas flow 12 L/min. The metabolites of interest were detected using a dynamic MRM mode. The MassHunter 10.0 software (Agilent Technologies) was used for the data processing. Ten-point linear calibration curves with internal standardization was constructed for the quantification of metabolites. Conditions were compared using Welch’s t-test, p-value was subsequently corrected for multiple testing according to the Benjamini and Hochberg procedure^43^. Pathway enrichment was performed testing significantly affected metaboliltes (p < 0.05) against the Small Molecule Pathway Database (SMPDB)^49^ with MetaboAnalyst^50^.

### Confocal microscopy

For the confocal imaging of HAP1 cells cells, high precision microscope cover glasses (Marienfeld) were coated with poly-L-lysine hydrobromide (p6282, Sigma-Aldrich) according to the manufacturer’s protocol. Cells were seeded onto cover glasses in normal growth medium and fixed in 4% formaldehyde solution (AppliChem) in PBS 1x after 24h of incubation. Permeabilization and blocking of samples was performed in blocking solution (5% FCS, 0.3% Triton X-100 in PBS 1x) in PBS 1x) for 1h rocking. Anti-HA Tag (Roche #11867423001) and anti-AIF (CST #5318) primary antibodies were diluted 1:500 in antibody dilution buffer (1% BSA, 0.3% Triton X-100 in PBS1x) and applied for 2h at room temperature, rocking. Samples were washed three times in antibody dilution buffer and anti-rat Alexa Fluor 488 (Thermo Fischer Scientific #A11006) and anti-rabbit Alexa Fluor 594 (Thermo Fischer Scientific #A11012) secondary antibodies were applied 1:500 in antibody dilution buffer for 1h at room temperature, rocking. After three times washing in antibody dilution buffer, nuclei were counterstained with DAPI 1:1000 in PBS 1x, for 10min, rocking. Cover glasses were mounted onto microscopy slides using ProLong Gold (Thermo Fischer Scientific #P36934) antifade mountant. Image acquisition was performed on a confocal laser scanning microscope (Zeiss LSM 780, Carl Zeiss AG), equipped with an Airyscan detector using ZEN black 2.3 (Carl Zeiss AG).

### NAD/NADH and ATP measurements

For luminescence-based assays, 10.000 HAP1 cells/well were plated in a 96-well plate in triplicates. After 6h, ATP and NAD(H) levels were measured by CellTiterGlo and NAD/NADH-Glo assays (Promega). Readings were normalized to cell numbers measured with Casy (OMNI Life Sciences) on a mirror plate.

### Data analysis and visualization

Exploratory data analysis and visualizations were performed in R-project version 3.6.0 (47) with RStudio IDE version 1.2.1578, ggplot2 (3.3.0), dplyr (0.8.5), readr (1.3.1).

## Acknowledgments

We are grateful to the ProMet and Biomedical Sequencing (BSF) facilities at CeMM/MUW. We thank the members of the Superti-Furga and Menche groups for critical discussions.

## Author contributions

E.G.: conceptualization, methodology, software, formal analysis, investigation, validation, visualization, project administration, writing; G.F., investigation, validation; U.G.: methodology, software, data curation, formal analysis, visualization; C.S.: software, formal analysis, visualization; F.M.: software, formal analysis, visualization; S.L.: investigation, validation; V.S. data curation, formal analysis, visualization; I.S.: investigation; B.A.: formal analysis, investigation; F.K.: investigation; E.M. software; A.S.: investigation; M.R.: conceptualization; A.B.: formal analysis, funding acquisition; J.M.: conceptualization, funding acquisition; G.S-F. conceptualization, project administration, funding acquisition, writing.

## Data and materials availability

Transcriptomics and metabolomics dataset will be deposited in public repositories. The complete genetic interaction dataset will be made available at http://sigil.cemm.at.

